# Imaging proteins sensitive to direct fusions using transient peptide-peptide interactions

**DOI:** 10.1101/2023.07.01.547312

**Authors:** Zoe Gidden, Curran Oi, Emily Johnston, Haresh Bhaskar, Susan Rosser, Simon G. J. Mochrie, Mathew H. Horrocks, Lynne Regan

## Abstract

Fluorescence microscopy enables specific visualization of proteins in living cells and has played an important role in our understanding of protein subcellular location and function. Some proteins, however, show altered localization and/or function when labeled using direct fusions to fluorescent proteins, making them difficult to study in live cells. Additionally, the resolution of fluorescence microscopy is limited to ∼200 nm, which is two orders of magnitude larger than the size of most proteins. To circumvent these challenges, we previously developed LIVE-PAINT, a live-cell super-resolution approach that takes advantage of short interacting peptides to transiently bind a fluorescent protein to the protein-of-interest. Here, we successfully use LIVE-PAINT to image yeast membrane proteins that do not tolerate the direct fusion of a fluorescent protein by using peptide tags as short as 5-residues. We also demonstrate that it is possible to resolve multiple proteins at the nanoscale concurrently using orthogonal peptide interaction pairs.

**FOR TABLE OF CONTENTS ONLY:** 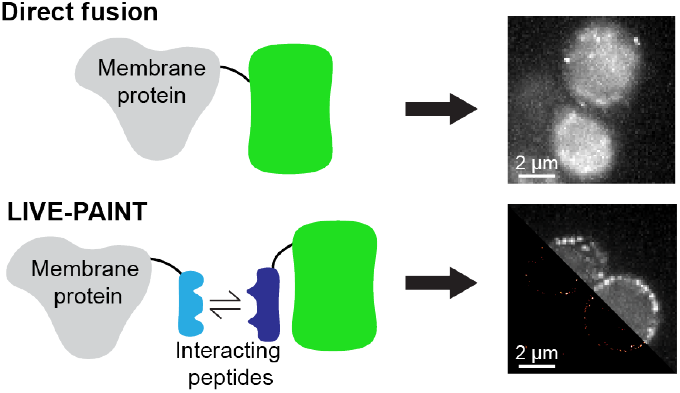

## INTRODUCTION

The ability to visualize proteins in their native cellular environment using a direct genetic fusion to a fluorescent protein (FP) has revolutionized cell biology. Unfortunately, not all proteins tolerate fusion to a FP, and can mislocalize or malfunction.^1,2^ One class of proteins that are frequently perturbed are yeast membrane transporter proteins, which show little or no localization to the plasma membrane when green fluorescent protein (GFP) is directly fused to their C-terminus.^3,4^ The highly abundant proton pump Pma1, for example, is well known to localize to the plasma membrane;^5–8^ however, Pma1 tagged with GFP at the C-terminus primarily localizes to the vacuole.^3,4^ Another easily perturbed protein in yeast is the nicotinic acid transporter Tna1, which is retained in the endoplasmic reticulum rather than localizing to the plasma membrane when tagged at the C-terminus with GFP.^9^

Imaging protein subcellular localization using fusions to FPs or other fluorescence techniques also has the drawback that the resolution is restricted to ∼200 nm due to the diffraction of light, unless a super-resolution (SR) technique is used.^10–12^ SR microscopy is an umbrella term for techniques that either illuminate a region of the sample smaller than the diffraction limit or use stochastic activation of fluorophores to enable the identification and precise localization of single emitters.^13^ Most SR techniques are challenging to apply to live-cell imaging, however, stimulated emission depletion (STED),^11^ reversible saturable optical fluorescence transition (RESOLFT)^14^; structured illumination microscopy (SIM)^15^; and some single-molecule localization microscopy (SMLM) approaches, such as Photoactivated Localization Microscopy (PALM)^10^, have been used to obtain SR images of proteins in live cells. For a comprehensive overview of live-cell SR techniques, see Godin, Lounis, and Cognet, 2014.^16^ Generally when applied to live-cell imaging, all of these techniques use FPs to label the protein being studied, which can be detrimental to normal localization and function. Furthermore, SR imaging of more than two proteins in live cells remains challenging unless SIM is used, but the improvement in resolution relative to diffraction limited imaging is only around two fold.^17^

We have recently developed a live-cell SR imaging method that can be applied to proteins that do not tolerate a direct fusion to a FP. Rather than directly fusing the protein-of-interest to a FP (∼25 kDa in size), <u>L</u>ive cell <u>I</u>maging using re<u>V</u>ersible int<u>E</u>ractions <u>P</u>oint <u>A</u>ccumulation for <u>I</u>maging in <u>N</u>anoscale <u>T</u>opography (LIVE-PAINT) uses non-covalent transient interactions between a peptide fused to the target protein, and the binding partner of the peptide fused to a FP.^18^ The use of the small peptide tag (< 5 kDa) renders this approach less perturbative than direct FP fusions. Here, we demonstrate that LIVE-PAINT can be used to locate a variety of difficult-to-image membrane proteins in yeast with nanometer precision. For proteins that are particularly sensitive to modifications, we show that it is possible to implement LIVE-PAINT on proteins tagged with only a 5-residue peptide (< 1 kDa).

Furthermore, using multiple orthogonal peptide-peptide interaction pairs and FPs with different emission wavelengths, we image two membrane-associated proteins simultaneously with nanometer precision. Although we demonstrate this functionality using membrane-associated proteins here, we expect LIVE-PAINT will enable us to visualize any difficult-to-label protein at the nanometer length scale. Additionally, LIVE-PAINT can be performed using any bright FP, unlike PALM, which requires photoactivatable or photoconvertible FPs.^12^ This means LIVE-PAINT has access to a much wider larger array of FPs, with varied absorption and emission spectra; this makes LIVE-PAINT an ideal SR method for tagging and imaging multiple target proteins concurrently.

## RESULTS

### Small peptide tags enable visualization of fusion-sensitive membrane proteins

Rather than relying on the genetically encoded fusion of full-length FPs to target proteins, LIVE-PAINT uses a peptide-peptide interaction pair to non-covalently and transiently associate a FP with the protein-of-interest (Figure 1ai, bi). We hypothesized that the fusion of a small peptide tag to the target protein would be less perturbative to the localization and/or function of the protein than direct fusion to a FP, and we therefore sought to apply LIVE-PAINT to image membrane proteins that mislocalize or have proven difficult to visualize when directly fused to GFP.

**Figure 1.**
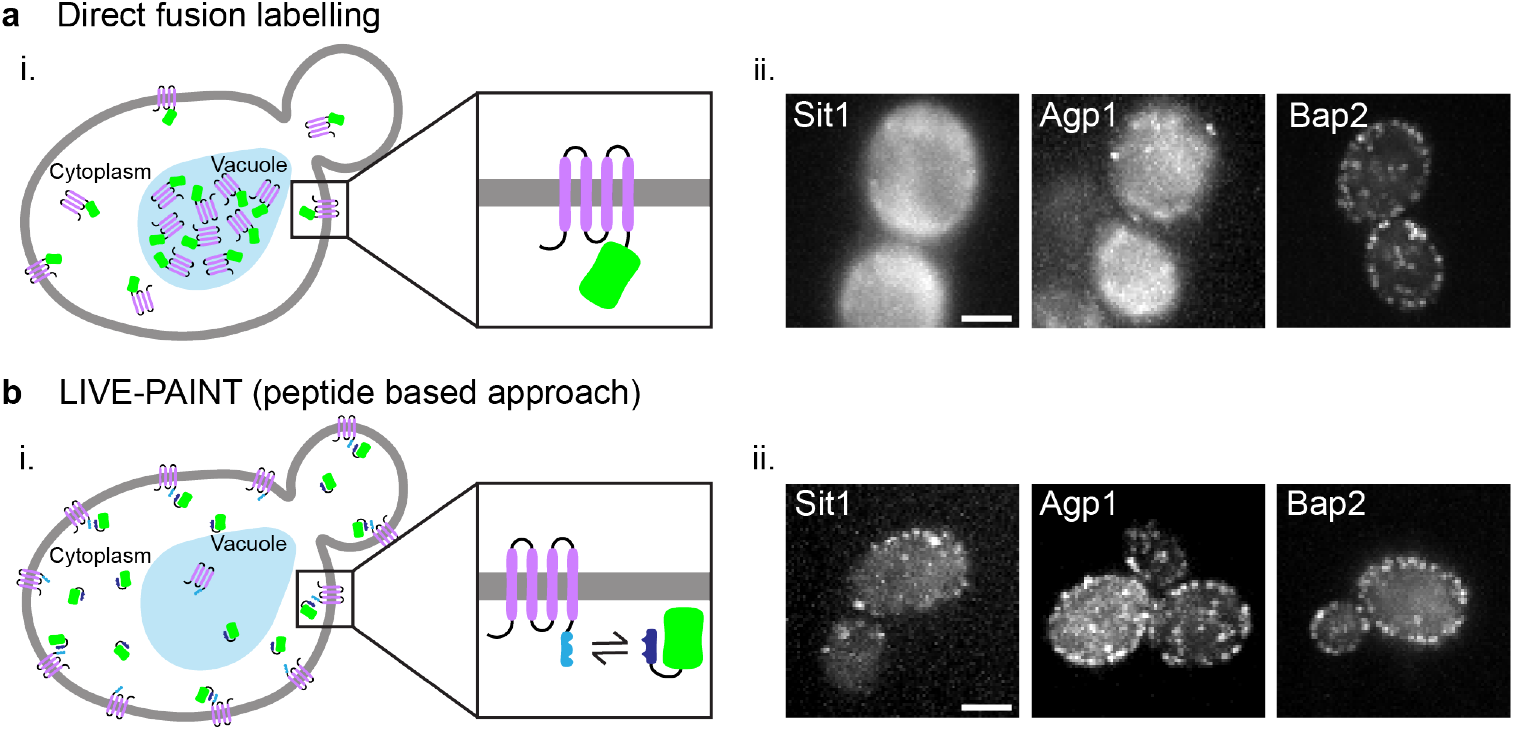
Membrane transporter proteins tolerate peptide fusions better than direct fusion to a FP. (ai) Membrane protein (purple) directly fused to a FP (green) often mislocalize to the vacuole (light blue oblong shape), or cytoplasm rather than locating to the plasma membrane (gray). (aii) Diffraction-limited TIRF images of three different membrane transporter proteins labelled by directly fusing mNG to their C-terminus. Scale bar is 2 μm and all images have the same dimensions. (bi) Membrane proteins fused to one half of a coiled-coil peptide (light blue rod) are less likely to mislocalize and can be imaged using the other half of the coiled-coil peptide (dark blue rod) fused to a FP. (bii) Diffraction-limited TIRF images the same three membrane transporter proteins tagged using the 101A/101B coiled coil pair: one half is fused to the C-terminus of the membrane protein and the other half is fused to mNG and expressed *in vivo*. Scale bar is 2 μm and all images have the same dimensions.

We first selected a set of *Saccharomyces cerevisiae* membrane transporter proteins that either accumulate at the vacuole or cannot be visualized upon direct fusion to GFP.^4^ Other researchers have previously carried out diffraction limited imaging of membrane proteins by labelling the target protein with a coiled-coil oriented outside of the cell and introducing the partner peptide in the imaging buffer so we anticipated our LIVE-PAINT approach would be feasible.^19^ We fused the coiled-coil peptide 101B to the C-terminus of each of these membrane proteins; the C-terminus for each of these proteins is predicted to be cytoplasmic.^20^ We also integrated a gene encoding the coiled-coil peptide 101A fused to mNeonGreen (mNG) into the genome driven by the galactose-inducible promoter pGAL1,^18^ replacing the *GAL2* gene in the process. The 101B peptide interacts with the 101A peptide with an estimated K_d_ of ∼200 nM.^21^

When imaged using total internal reflection fluorescence (TIRF) microscopy, membrane proteins tagged using the LIVE-PAINT system appear as a ring around the periphery of the cell. However, this does depend on the orientation of the yeast on the slide, the TIR angle, and the z plane used for imaging. Figure 1 shows three membrane proteins, Sit1, Agp1, and Bap2, imaged either using a direct fusion (Figure 1aii) to mNG or using a peptide-peptide interaction pair to associate mNG to the target protein (Figure 1bii). We observe that when Sit1 is directly fused to mNG it does not localize to the membrane, whereas for Agp1 and Bap2, direct fusion to mNG only partially affects their membrane localization. All three proteins show improved membrane localization when tagged with 101B and imaged with 101A-mNG.

We subsequently used LIVE-PAINT to image a collection of 12 plasma membrane proteins that have been reported to exhibit partial or complete mislocalization when directly fused to GFP in yeast (Figure 2a).^4^ For the negative control, 101A-mNG expressed in yeast in the absence of a target protein fused to 101B, we did not observe more than background fluorescence at any specific location within the cell including at the plasma membrane (Figure S1). We found that the success of this approach does not depend on the abundance of the membrane protein, although there is generally improved contrast between membrane signal and background signal for more abundant proteins (see Table S1 for approximate abundance and Figure 2a). We believe that this effect is because the fraction of FP bound to the target protein should increase as the concentration of target protein increases.

**Figure 2.**
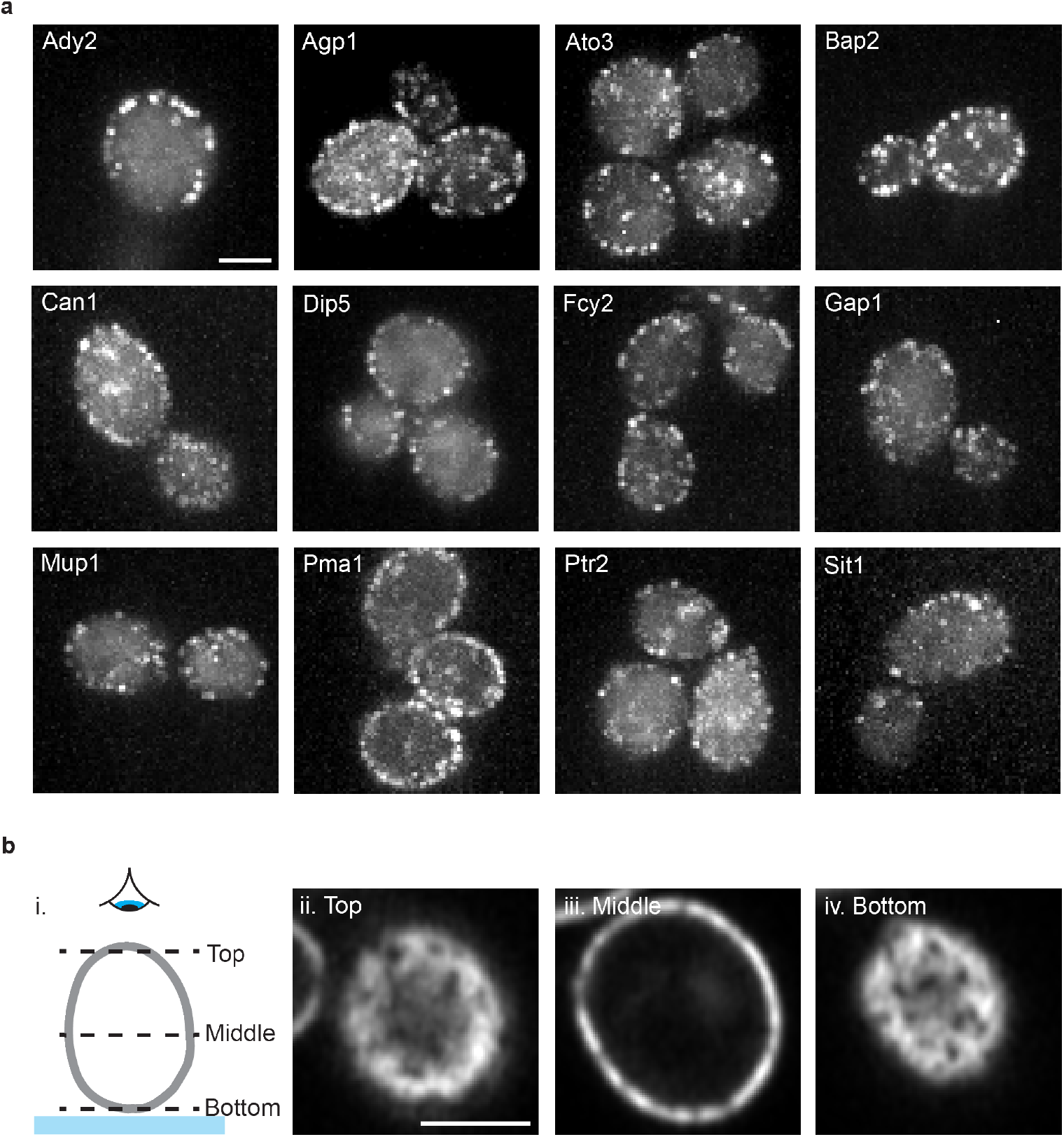
Membrane proteins imaged using the peptide tagging approach show fluorescent signal at the plasma membrane in live cells. (a) Diffraction-limited TIRF images of membrane transporter proteins tagged with the 101B peptide at their C-terminus and imaged by co-expressing 101A-mNG. Scale bar in (a) is 2 μm and all images have the same dimensions. (bi) Schematic representation of z slices through a yeast cell shown in (bii) - (biv). Glass slide represented by the blue band and an eye showing the view from above. (bii) - (biv) Top, middle, and bottom z slices of Pma1-101B imaged by co-expressing 101A-mNG acquired using an Airyscan microscope. Scale bar in (bii) is 2 μm and all images (bii) - (biv) have the same dimensions.

We have also used LIVE-PAINT to image Pma1 at different planes through the yeast cell using a Zeiss LSM880 confocal microscope with Airyscan (Figure 2b) enabling a 3D rendering of distribution throughout the entire cell (Supplementary video 1). Clear plasma membrane signal with minimal internal signal is observed in the plane that cuts through the middle of the cell (Figure 2biii). In both planes that cut through the membrane at the top and bottom of the cell, regions where Pma1 is excluded can be clearly observed (Figure 2bii & 2biv). This is consistent with the network-like distribution, exclusive of membrane compartment of Can1 domains or eisosomes, which has previously been described for Pma1. This is expected as Pma1 is known to be arranged in microcompartments at the membrane.^22^

### Super-resolution imaging of membrane proteins reveals closely spaced protein clusters

We next selected four membrane transporter proteins that showed clear localization at the plasma membrane when tagged using 101A/101B for super-resolution imaging using LIVE-PAINT (Figure 3a). In less than two minutes of data acquisition for Pma1, LIVE-PAINT led to 367 +/-315 localizations (mean +/-SD, n = 15 cells) with a precision of 10.7 +/-0.4 nm (mean +/-SD, n = 15 cells), leading to a resolution of 67.3 +/-13.4 nm (mean +/-SD, n = 15 cells) (calculated using Fourier Ring Correlation (10 Brink, T. RustFRC [Computer software]) (see Table S2 for a summary of resolution, precision, and number of localizations achieved for each protein presented in Figure 3). The success of the labelling strategy was measured by quantifying the percentage of localizations at the membrane (Figure 3b). We found that 60%, 70%, 67% and 82% of total localizations were at the membrane for Ato3, Bap2, Dip5, and Pma1, respectively. The resulting images revealed clusters of localizations spaced closer than 200 nm apart (Figure 3c). These clusters are spaced too close to one another to be distinguished by diffraction-limited microscopy; this highlights the importance of imaging such proteins using SR methods.

**Figure 3.**
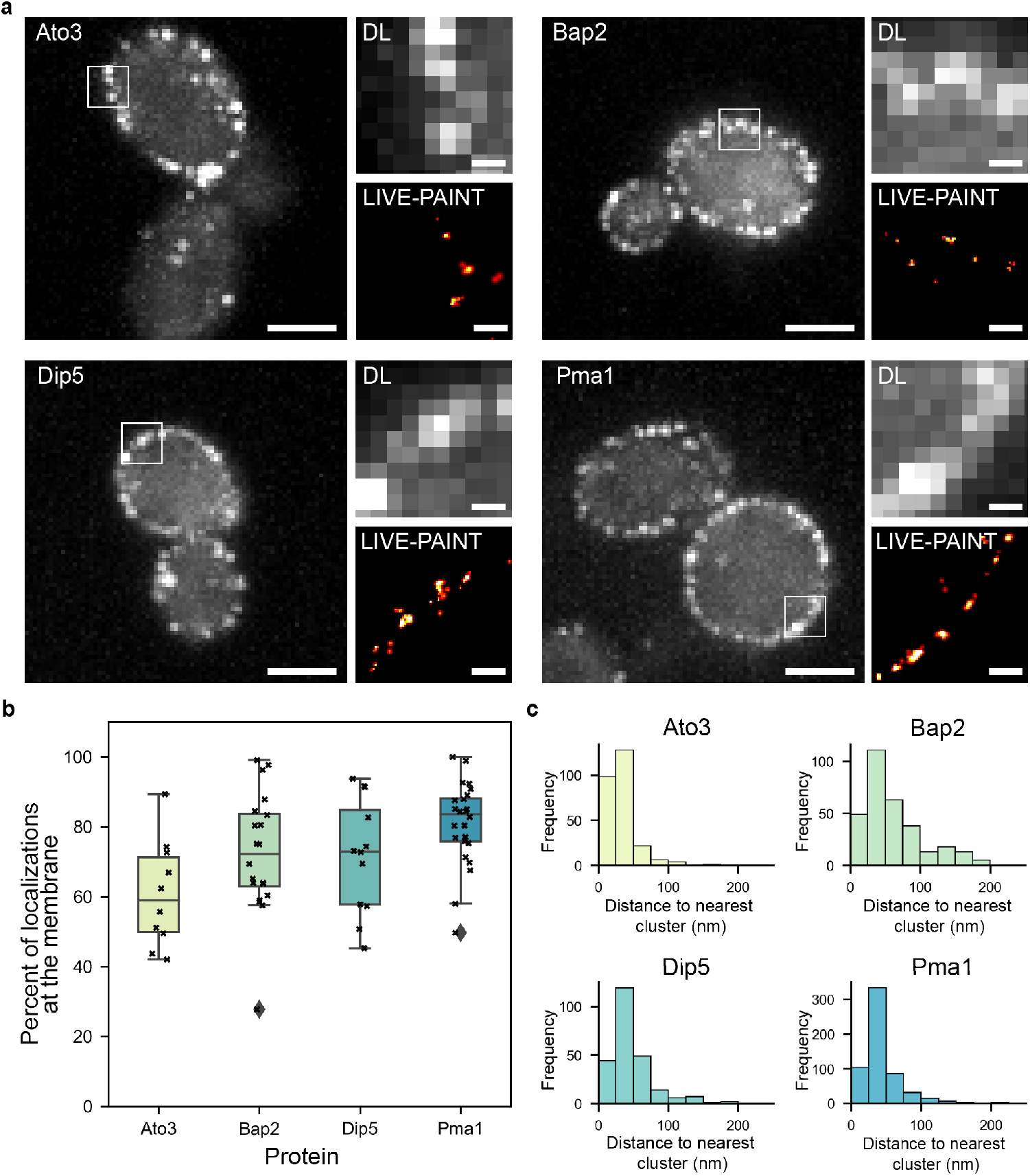
Live cell imaging of membrane proteins using LIVE-PAINT measures protein localization with nanometer precision. (a) Diffraction-limited and super-resolution LIVE-PAINT images for four representative membrane transporter proteins; Ato3, Bap2, Dip5, and Pma1. For these proteins, 101B is fused to the C-terminus and imaged by co-expressing 101A-mNG. Scale bars are 2 μm for full-cell images and 250 nm for zoom ins. Full-cell LIVE-PAINT images are shown in Figure S2. (b) Box plots showing the percentage of total localizations at the membrane for each of the membrane proteins. Classification of membrane localizations is shown in Figure S3. (c) Histograms of the distance to the nearest cluster for each of the membrane proteins. Ato3 (n = 4 cells), Bap2 (n = 15 cells), Dip5 (n = 6 cells), and Pma1 (n = 13 cells).

### A five-residue fusion tag is sufficient to enable live-cell super-resolution imaging

Although most proteins will tolerate fusion to a 5 kDa peptide tag, sometimes this may be too large, and we therefore sought to demonstrate LIVE-PAINT with a shorter peptide tag. For this purpose, we selected the 5-residue KQTSV peptide that binds reversibly to the 11 kDa protein PDZ3. To test this system, we fused KQTSV to the endogenous septum protein Cdc12 and co-expressed PDZ3 protein fused to mNG under the galactose inducible promoter (Figure S4). While it was possible to observe some fluorescence at the septum (diffraction-limited and LIVE-PAINT images shown in Figure S4b), there was also significant background fluorescence in the cell, and the spatial resolution was lower than expected (269 +/-108 nm, mean +/-SD, n = 5 cells). We reasoned that this was due to the low affinity of the KQTSV/PDZ3 system (K_D_ ∼800 nM),^23^ and we therefore trialed tagging mNG with two tandem repeats of the PDZ3 protein, in an approach analogous to that utilized with DNA-PAINT to enhance the signal-to-background ratio.^24^ As expected, this led to clearer images of the septum (Figure S4b), and a higher resolution was obtained (123 +/-37 nm, mean +/-SD, n = 14 cells) (imaging of Cdc12 using 101A/B and the negative control with 2xPDZ3 only is also shown in Figure S4).

After establishing the feasibility of using the 2xPDZ3/KQTSV protein-peptide pair for LIVE-PAINT imaging, we used this interaction pair to image two more proteins: Pma1, which is not amenable to direct fusions to GFP, and Pil1, which is a membrane-associated protein (Figure 4). Similar to the 101A/B peptide pair, imaging using the 2xPDZ3/KQTSV interaction pair produces clear plasma membrane localizations for both Pma1 and Pil1 with very little internal signal for both the diffraction-limited and SR images (Figure 4b and c). To quantify the success of the labeling strategy, the percentage of membrane specific to total localizations was calculated for each protein (Figure 4d). On average, for Pma1, 83% of localizations were at the membrane and for Pil1 74% of localizations were membrane specific. See Table S3 for a summary of resolution, precision, and number of localizations achieved for Pma1 and Pil1 imaged with the 2xPDZ3/KQTSV protein-peptide pair. The success of this labeling approach demonstrates the generalizability of LIVE-PAINT: other interaction pairs, not only 101A/101B, can be used to achieve clear labeling of membrane proteins that mislocalize when directly fused to GFP.

**Figure 4.**
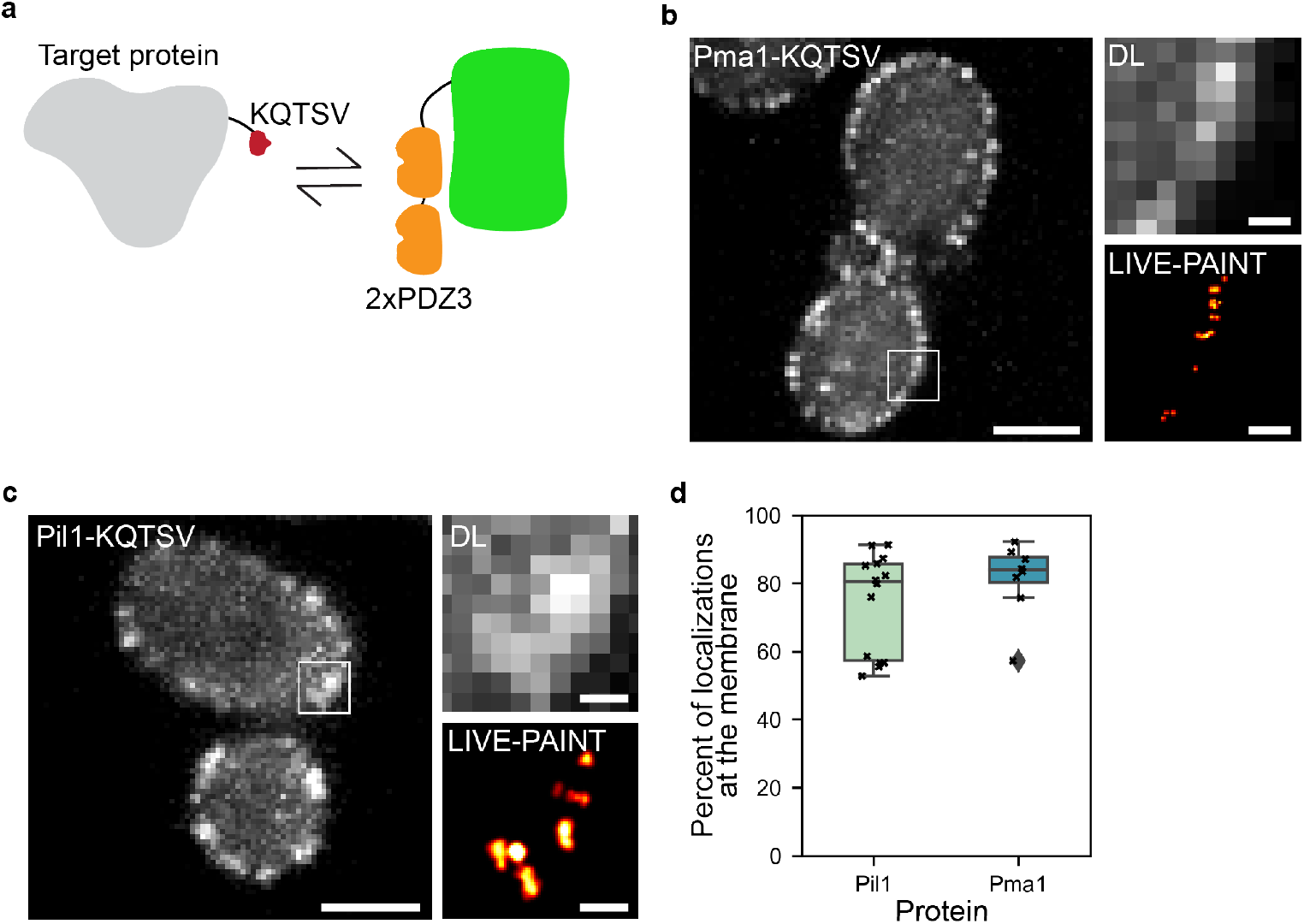
The 5-residue tag KQTSV can be used for LIVE-PAINT imagining of membrane proteins in live cells. (a) Schematic representation of the labelling strategy. The short KQTSV peptide (red) is used to label the target protein, and this reversibly binds to the 2xPDZ3 protein (orange units) which are attached to mNG. (b)-(c) Diffraction-limited (DL) and super-resolution LIVE-PAINT images for two membrane associated proteins. For these proteins, KQTSV is fused to the C-terminus and imaged by co-expressing 2xPDZ3-mNG. Scale bars are 2 μm for full-cell images and 250 nm for zoom ins. Full-cell LIVE-PAINT images are shown in Figure S5. (d) Box plots showing the percentage of total localizations at the membrane for Pil1 and Pma1. Classification of membrane localizations is shown in Figure S6.

### LIVE-PAINT enables simultaneous live-cell super-resolution imaging of two proteins

As two-color live-cell SR imaging is challenging with current methods and often requires a direct fusion to a FP, we sought to use LIVE-PAINT to image two proteins simultaneously in live cells. We chose to image two plasma membrane associated proteins, Arc35 and Pil1, which are predicted to be close together but not localized to the same structures. Arc35 is a component of actin patches and assists in the organization of actin to facilitate endocytosis^25^ and Pil1 is a BAR domain protein that facilitates the formation of eisosome subdomains of the plasma membrane,^26^ which are associated with sites of protection from endocytosis.^27^ We used two peptide interaction pairs, 101A/101B and 108A/108B, that have previously been shown to be orthogonal,^21^ to C-terminally tag Arc35 and Pil1, respectively (Figure 5a).^21^ 101A fused to mNG was integrated into the genome and expressed under the galactose inducible promoter, pGAL1, as before, and 108A fused to mCherry was also integrated into the genome and expressed under the same promoter.

**Figure 5.**
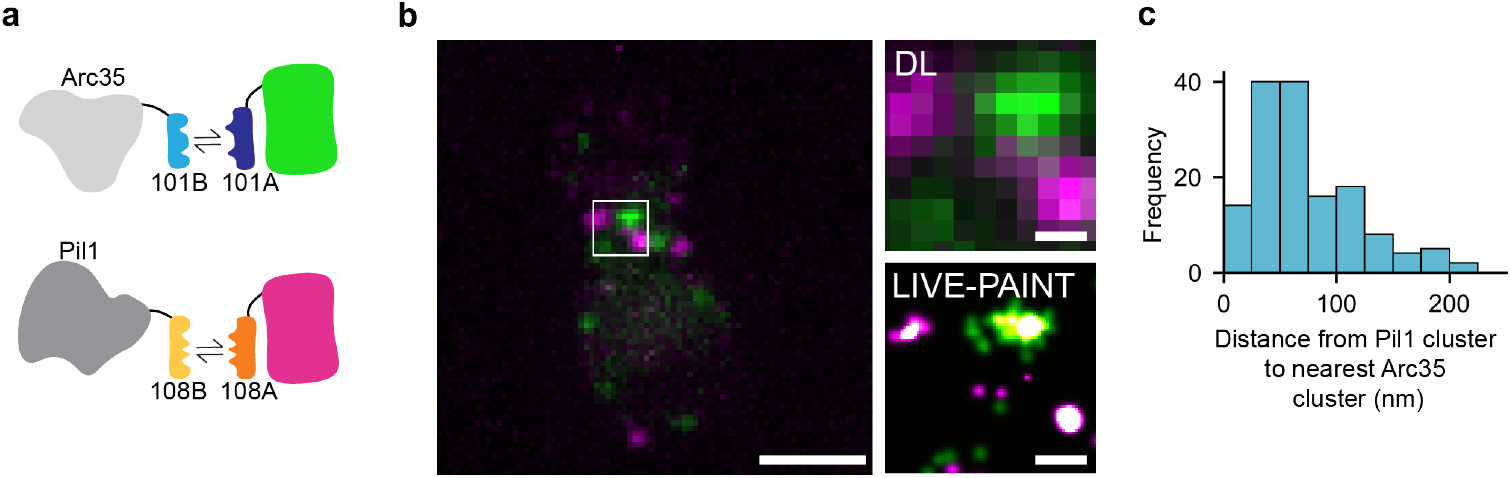
LIVE-PAINT can be used to image two proteins in live cells, concurrently, with nanometer precision. (a) Schematic representation of the LIVE-PAINT labelling strategy used to image Arc35 and Pil1 simultaneously. The orthogonal peptide pairs 101A/101B and 108A/108B were used to label Arc35 and Pil1 with mNG and mCherry respectively. (b) Diffraction-limited and SR images of Arc35 (green) and Pil1 (magenta) simultaneously imaged using the LIVE-PAINT. Scale bars are 2 μm for ful-cell images and 250 nm for zoom ins. (c) Histogram showing the distance between each Pil1 cluster and its closest Arc35 cluster in the same cell (n = 6 cells).

The SR images generated show that Arc35 and Pil1 are arranged in clusters with little to no overlap between the two proteins (Figure 5b). We achieved a resolution of 83 +/-22 nm (mean +/-SD, n = 6 cells) with mNG and 68 +/-24 nm (mean +/-SD, n = 6 cells) with mCherry (10 Brink, T. RustFRC [Computer software]) (see Table S4 for a summary of resolution, precision and number of localizations achieved for Arc35 and Pil1 in Figure 4). For each Pil1 cluster, we calculated the distance to the nearest Arc35 cluster (Figure 5c) and found that all the nearest clusters were closer than the diffraction-limit of light (210 nm), meaning that they would not be spatially distinguished using standard fluorescence microscopy. This demonstrates the value of using a live-cell SR imaging technique for concurrent imaging of proteins such as Arc35 and Pil1 that localize near each other in the cell.

## CONCLUSIONS AND DISCUSSION

We have demonstrated that proteins sensitive to direct fusion to GFP can be fused to a small peptide and imaged in live cells using the binding partner of the peptide fused to a FP. We used membrane transporter proteins as an example class of proteins that are generally sensitive to direct fusion to GFP and show that they generally tolerate fusion to a small (< 5 kDa) peptide and subsequent LIVE-PAINT imaging. Our approach clearly recovers the expected localization of the tagged protein in 12 membrane transporter proteins we tagged and imaged. We also carried out 3D imaging on one of the membrane proteins, Pma1, using a peptide tagging approach. Additionally, we have also demonstrated that we can perform LIVE-PAINT SR imaging of multiple proteins in yeast, both separately and simultaneously using two orthogonal peptide-peptide interaction pairs.

We expect this approach to be broadly useful for tagging and imaging proteins sensitive to direct fusions to a FP. Here, we have demonstrated the use of LIVE-PAINT for imaging yeast membrane transporter proteins, but the small size of the peptides used relative to a FP suggests that this approach will be useful for visualizing other difficult-to-label proteins.^4,28,29^ We note that while we successfully used LIVE-PAINT to image membrane proteins with a variety of abundances, we generally found that higher abundance proteins produced images with clearer membrane signal.

In addition to showing that our peptide tagging approach can be less perturbative to the localization of the target protein, we showed that the binding affinity of the interaction peptide-peptide or peptide-protein pairs was important for successful LIVE-PAINT imaging. Through this work and our previous work, we have found that interaction pairs with a binding affinity between 1-300 nM work well^18^; however, here we also show that weaker binding pairs can potentially be used if the number of binding sites available on the imaging strand is increased. This enables the use of shorter, less perturbative peptides.

Finally, we demonstrate that LIVE-PAINT can be used to image two targets simultaneously in live cells. This approach could be used to carry out live-cell colocalization studies on proteins that do not tolerate direct fusions and to investigate colocalization at resolutions higher than the diffraction-limit of light. Two-color imaging using E/K coiled-coil interaction pairs has also been demonstrated by Eklund & Jungmann in fixed mammalian cells.^30^ To date, we have used 5 different interaction pairs for LIVE-PAINT imaging: the protein-peptide interaction pairs TRAP4/MEEVF and 2xPDZ3/KQTSV, and the peptide-peptide interaction pairs SYNZIP17/SYNZIP18, 101A/B and 108A/B.^18^ The diversity of interaction pairs suitable for LIVE-PAINT illustrates the broad potential for this approach for tagging and imaging proteins sensitive to direct fusions to FPs. Similarly, the growing set of orthogonal interaction pairs we have shown to be suitable for LIVE-PAINT reveals the potential for simultaneous live-cell SR imaging of multiple proteins; here, we demonstrate this with two proteins, but we envision that simultaneous LIVE-PAINT imaging of three or more proteins is also possible.

## ASSOCIATED CONTENT

### Supporting Information

Materials and methods; negative control for LIVE-PAINT imaging with 101A-mNeonGreen and 2xPDZ3-mNeonGreen (Figure S1 and S4 respectively); comparison of LIVE-PAINT imaging with 101A, 1xPDZ3, or 2xPDZ3 fused to mNeonGreen (Figure S4); full-cell super-resolution images of proteins imaged using LIVE-PAINT (Figure S2 and S5); representative plots to show localization classification as membrane, cellular or external (Figure S3 and S6); list of proteins imaged in this study (Table S1); summary statistics for all super-resolution images (Table S2-S4); list of primers used in this study (Table S5 and S6), list of yeast strains used in this study (Table S7) (PDF).

Movie of a 3D rendering of Pma1 labelled using LIVE-PAINT (Video S1) (AVI)

## AUTHOR INFORMATION

### Author Contributions

C.O., Z.G., M.H.H., and L.R. designed experiments. AiryScan imaging was carried out by E.J. Z.G. and C.O. performed all other experiments. Z.G., C.O., M.H.H., H.B., E.J., S.R., S.G.J.M., and L.R. analyzed data. The manuscript was written by Z.G., C.O., M.H.H., and L.R., with contributions from all authors. All authors have given approval to the final version of the manuscript.

### Notes

The authors declare no competing financial interests.

## ACKNOWLEDGMENT

Confocal microscopy imaging was performed with Dr Toni McHugh at Centre Optical Instrumentation Laboratory (COIL), which is supported by a Core Grant (203149) to the Wellcome Centre for Cell Biology at the University of Edinburgh. We would also like to thank Dr Katherine Paine, Dr Ella Thornton, Dr Raef Shams, Dr Mai-Britt Jensen, Rossana Boni, Noelia Pelegrina-Hidalgo, Kasia Stafaniak, and Sneha Ravi, for thoughtful comments on the manuscript.

## ABBREVIATIONS

FP: fluorescent protein
GFP: green fluorescent protein
SR: super-resolution
STED: stimulated emission depletion
RESOLFT: reversible saturable optical fluorescence transition
SIM: structured illumination microscopy
SMLM: single-molecule localization microscopy
PALM: photoactivation localization microscopy
LIVE-PAINT: Live cell imaging using reversible interactions point accumulation for imaging in nanoscale topography
mNG: mNeonGreen
TIRF: total internal reflection fluorescence
DL: diffraction-limited.

## REFERENCES

(1) Agbulut, O.; Coirault, C.; Niederländer, N.; Huet, A.; Vicart, P.; Hagège, A.; Puceat, M.; Menasché, P. GFP Expression in Muscle Cells Impairs Actin-Myosin Interactions: Implications for Cell Therapy. Nat. Methods 2006, 3 (5), 331–331. https://doi.org/10.1038/nmeth0506-331.

(2) Lisenbee, C. S.; Karnik, S. K.; Trelease, R. N. Overexpression and Mislocalization of a Tail-Anchored GFP Redefines the Identity of Peroxisomal ER. Traffic 2003, 4 (7), 491–501. https://doi.org/10.1034/j.1600-0854.2003.00107.x.

(3) Hinrichsen, M.; Lenz, M.; Edwards, J. M.; Miller, O. K.; Mochrie, S. G. J.; Swain, P. S.; Schwarz-Linek, U.; Regan, L. A New Method for Post-Translationally Labeling Proteins in Live Cells for Fluorescence Imaging and Tracking. Protein Eng. Des. Sel. PEDS 2017, 30 (12), 771–780. https://doi.org/10.1093/protein/gzx059.

(4) Huh, W.-K.; Falvo, J. V.; Gerke, L. C.; Carroll, A. S.; Howson, R. W.; Weissman, J. S.; O’Shea, E. K. Global Analysis of Protein Localization in Budding Yeast. Nature 2003, 425 (6959), 686–691. https://doi.org/10.1038/nature02026.

(5) Mason, A. B.; Allen, K. E.; Slayman, C. W. C-Terminal Truncations of the Saccharomyces Cerevisiae PMA1 H+-ATPase Have Major Impacts on Protein Conformation, Trafficking, Quality Control, and Function. Eukaryot. Cell 2014, 13 (1), 43–52. https://doi.org/10.1128/EC.00201-13.

(6) Rao, R.; Drummond-Barbosa, D.; Slayman, C. W. Transcriptional Regulation by Glucose of the Yeast PMA1 Gene Encoding the Plasma Membrane H+-ATPase. Yeast 1993, 9 (10), 1075–1084. https://doi.org/10.1002/yea.320091006.

(7) Serrano, R. Characterization of the Plasma Membrane ATPase of Saccharomyces Cerevisiae. Mol. Cell. Biochem. 1978, 22 (1), 51–63. https://doi.org/10.1007/BF00241470.

(8) Szopinska, A.; Degand, H.; Hochstenbach, J.-F.; Nader, J.; Morsomme, P. Rapid Response of the Yeast Plasma Membrane Proteome to Salt Stress. Mol. Cell. Proteomics MCP 2011, 10 (11), M111.009589. https://doi.org/10.1074/mcp.M111.009589.

(9) Paine, K. M.; Ecclestone, G. B.; MacDonald, C. Fur4-Mediated Uracil-Scavenging to Screen for Surface Protein Regulators. Traffic 2021, 22 (11), 397–408. https://doi.org/10.1111/tra.12815.

(10) Betzig, E.; Patterson, G. H.; Sougrat, R.; Lindwasser, O. W.; Olenych, S.; Bonifacino, J. S.; Davidson, M. W.; Lippincott-Schwartz, J.; Hess, H. F. Imaging Intracellular Fluorescent Proteins at Nanometer Resolution. Science 2006, 313 (5793), 1642–1645. https://doi.org/10.1126/science.1127344.

(11) Hell, S. W.; Wichmann, J. Breaking the Diffraction Resolution Limit by Stimulated Emission: Stimulated-Emission-Depletion Fluorescence Microscopy. Opt. Lett. 1994, 19 (11), 780–782. https://doi.org/10.1364/ol.19.000780.

(12) Rust, M. J.; Bates, M.; Zhuang, X. Sub-Diffraction-Limit Imaging by Stochastic Optical Reconstruction Microscopy (STORM). Nat. Methods 2006, 3 (10), 793–796. https://doi.org/10.1038/nmeth929.

(13) Horrocks, M. H.; Palayret, M.; Klenerman, D.; Lee, S. F. The Changing Point-Spread Function: Single-Molecule-Based Super-Resolution Imaging. Histochem. Cell Biol. 2014, 141 (6), 577–585. https://doi.org/10.1007/s00418-014-1186-1.

(14) Hofmann, M.; Eggeling, C.; Jakobs, S.; Hell, S. W. Breaking the Diffraction Barrier in Fluorescence Microscopy at Low Light Intensities by Using Reversibly Photoswitchable Proteins. Proc. Natl. Acad. Sci. U. S. A. 2005, 102 (49), 17565–17569. https://doi.org/10.1073/pnas.0506010102.

(15) Heintzmann, R.; Cremer, C. G. Laterally Modulated Excitation Microscopy: Improvement of Resolution by Using a Diffraction Grating. In Optical Biopsies and Microscopic Techniques III; SPIE, 1999; Vol. 3568, pp 185–196. https://doi.org/10.1117/12.336833.

(16) Godin, A. G.; Lounis, B.; Cognet, L. Super-Resolution Microscopy Approaches for Live Cell Imaging. Biophys. J. 2014, 107 (8), 1777–1784. https://doi.org/10.1016/j.bpj.2014.08.028.

(17) Gustafsson, M. G. Surpassing the Lateral Resolution Limit by a Factor of Two Using Structured Illumination Microscopy. J. Microsc. 2000, 198 (Pt 2), 82–87. https://doi.org/10.1046/j.1365-2818.2000.00710.x.

(18) Oi, C.; Gidden, Z.; Holyoake, L.; Kantelberg, O.; Mochrie, S.; Horrocks, M. H.; Regan, L. LIVE-PAINT Allows Super-Resolution Microscopy inside Living Cells Using Reversible Peptide-Protein Interactions. Commun. Biol. 2020, 3 (1), 1–10. https://doi.org/10.1038/s42003-020-01188-6.

(19) Yano, Y.; Yano, A.; Oishi, S.; Sugimoto, Y.; Tsujimoto, G.; Fujii, N.; Matsuzaki, K. Coiled-Coil Tag—Probe System for Quick Labeling of Membrane Receptors in Living Cells. ACS Chem. Biol. 2008, 3 (6), 341–345. https://doi.org/10.1021/cb8000556.

(20) Möller, S.; Croning, M. D. R.; Apweiler, R. Evaluation of Methods for the Prediction of Membrane Spanning Regions. Bioinformatics 2001, 17 (7), 646–653. https://doi.org/10.1093/bioinformatics/17.7.646.

(21) Chen, R.; Rishi, H. S.; Potapov, V.; Yamada, M. R.; Yeh, V. J.; Chow, T.; Cheung, C. L.; Jones, A. T.; Johnson, T. D.; Keating, A. E.; DeLoache, W. C.; Dueber, J. E. A Barcoding Strategy Enabling Higher-Throughput Library Screening by Microscopy. ACS Synth. Biol. 2015, 4 (11), 1205–1216. https://doi.org/10.1021/acssynbio.5b00060.

(22) Malínská, K.; MalínskÝ, J.; Opekarová, M.; Tanner, W. Visualization of Protein Compartmentation within the Plasma Membrane of Living Yeast Cells. Mol. Biol. Cell 2003, 14 (11), 4427–4436. https://doi.org/10.1091/mbc.e03-04-0221.

(23) Gianni, S.; Engström, Å.; Larsson, M.; Calosci, N.; Malatesta, F.; Eklund, L.; Ngang, C. C.; Travaglini-Allocatelli, C.; Jemth, P. The Kinetics of PDZ Domain-Ligand Interactions and Implications for the Binding Mechanism*. J. Biol. Chem. 2005, 280 (41), 34805–34812. https://doi.org/10.1074/jbc.M506017200.

(24) Clowsley, A. H.; Kaufhold, W. T.; Lutz, T.; Meletiou, A.; Di Michele, L.; Soeller, C. Repeat DNA-PAINT Suppresses Background and Non-Specific Signals in Optical Nanoscopy. Nat. Commun. 2021, 12 (1), 501. https://doi.org/10.1038/s41467-020-20686-z.

(25) Schaerer-Brodbeck, C.; Riezman, H. Saccharomyces Cerevisiae Arc35p Works through Two Genetically Separable Calmodulin Functions to Regulate the Actin and Tubulin Cytoskeletons. J. Cell Sci. 2000, 113 (3), 521–532. https://doi.org/10.1242/jcs.113.3.521.

(26) Ziółkowska, N. E.; Karotki, L.; Rehman, M.; Huiskonen, J. T.; Walther, T. C. Eisosome-Driven Plasma Membrane Organization Is Mediated by BAR Domains. Nat. Struct. Mol. Biol. 2011, 18 (7), 854–856. https://doi.org/10.1038/nsmb.2080.

(27) Brach, T.; Specht, T.; Kaksonen, M. Reassessment of the Role of Plasma Membrane Domains in the Regulation of Vesicular Traffic in Yeast. J. Cell Sci. 2011, 124 (3), 328–337. https://doi.org/10.1242/jcs.078519.

(28) Montecinos-Franjola, F.; Bauer, B. L.; Mears, J. A.; Ramachandran, R. GFP Fluorescence Tagging Alters Dynamin-Related Protein 1 Oligomerization Dynamics and Creates Disassembly-Refractory Puncta to Mediate Mitochondrial Fission. Sci. Rep. 2020, 10 (1), 14777. https://doi.org/10.1038/s41598-020-71655-x.

(29) Meyer, T.; Begitt, A.; Vinkemeier, U. Green Fluorescent Protein-Tagging Reduces the Nucleocytoplasmic Shuttling Specifically of Unphosphorylated STAT1. FEBS J. 2007, 274 (3), 815–826. https://doi.org/10.1111/j.1742-4658.2006.05626.x.

(30) Eklund, A. S.; Jungmann, R. Optimized Coiled-Coil Interactions for Multiplexed Peptide-PAINT. Small 2023, 19 (12), 2206347. https://doi.org/10.1002/smll.202206347.

