## Supplementary Information for "Imaging proteins sensitive to direct fusions using transient peptide-peptide interactions"

[1]School of Biological Sciences, University of Edinburgh, Edinburgh, EH9 3DW, United Kingdom, [2]EaStCHEM School of Chemistry, The University of Edinburgh, Edinburgh, EH9 3FJ, United Kingdom, [3]Department of Genome Sciences, University of Washington, Seattle, WA 98195, United States, [4]Centre for Engineering Biology, University of Edinburgh, Edinburgh EH9 3BD, United Kingdom, [5]Department of Physics, Yale University, New Haven, CT 06520, United States, [6]Integrated Graduate Program in Physical and Engineering Biology, Yale University, New Haven, CT 06520, United States, [7]IRR Chemistry Hub, Institute for Regeneration and Repair, The University of Edinburgh, Edinburgh, EH16 4UU, United Kingdom, [8]Institute of Quantitative Biology, Biochemistry and Biotechnology, Edinburgh, EH9 3FF, United Kingdom

‡ Z.G. and C.O. contributed equally to this work


| Materials and Methods | |
|---|---|
| Figure S1 | 101A-mNeonGreen alone does not localize to the plasma membrane. |
| Figure S2 | Full-cell super-resolution images of membrane proteins labelled with the 101A/B peptide pair shown in Figure 3. |
| Figure S3 | Representative plots showing localizations classified as membrane, cellular or external for membrane proteins labelled with the 101A/B peptide pair. |
| Figure S4 | KQTSV can be used as a peptide label for LIVE-PAINT imaging with two tandem repeats of PDZ3 fused to mNeonGreen. |
| Figure S5 | Full-cell super-resolution images of membrane proteins labelled with the KQTSV/2xPDZ3 peptide-protein pair shown in Figure 4. |
| Figure S6 | Representative plots showing localizations classified as membrane, cellular or external for membrane proteins labelled with the KQTSV/2xPDZ3 peptide-protein pair. |
| Table S1 | Membrane transporter proteins tagged and imaged in this study. |
| Table S2 | Summary statistics of super-resolution images of membrane associated proteins imaged using LIVE-PAINT in Figure 3. |
| Table S3 | Summary statistics of super-resolution images of membrane associated proteins imaged using LIVE-PAINT in Figure 4. |
| Table S4 | Summary statistics of super-resolution images of Arc35 and Pil1 imaged using two color LIVE-PAINT and shown in Figure 5. |
| Table S5 | Primers for tagging membrane proteins at their genomic loci. |
| Table S6 | Primers for generating a yeast strain with two membrane-associated proteins tagged at their genomic loci. |
| Table S7 | List of yeast strains used in this study. |
| References | |

MATERIALS AND METHODS

### Molecular biology

All cloning was performed in TOP10 *E. coli*, using standard techniques. All constructs were generated in the pFA6a-His3MX6 and pFA6a-KanMX6 yeast integration vectors. All linker sequences used are GGSGSGLQ. Plasmids were constructed via Gibson assembly using NEBuilder® HiFi DNA Assembly Master Mix (New England Biolabs) by amplifying the plasmid backbone via PCR and inserting other parts of the construct either using gBlocks (Integrated DNA Technologies) or other PCR amplified sequences.

### Yeast strain construction

Except where otherwise noted, standard methods for genetically modifying yeast and preparing growth media were used.[1]

To label membrane transporter proteins we used the 101A/101B coiled coil interaction pair,[2] which we have previously demonstrated to be compatible with LIVE-PAINT imaging.[3] At the gene level, 101B was fused to the C-terminus of the target membrane protein at its endogenous locus. 101A was fused to mNG, under control of the galactose-inducible promoter, and integrated into the genome, replacing *GAL2*. When *GAL2* is deleted, the galactose-inducible promoter has a linear response with increasing galactose concentration.[4] This same approach was used when tagging Pma1 using the KQTSV/PDZ3 interaction pair.

Yeast strains were produced by amplifying the insert and selection marker from a yeast integration plasmid (pFA6a-His3MX6 or pFA6a-KanMX6) with ~45 bp overhang sequences matching the 45 bp upstream of the target protein's stop codon and ~45 bp downstream of the desired integration site in the genome. The primers used to tag the various proteins studied in this work are listed in Supplementary Table 1. To transform the yeast, a 3 mL culture of yeast was grown overnight in YPD, back-diluted to an $OD_{600}$ of 0.1 in 3 mL in the morning and grown to $OD_{600}$ of ~0.6-0.8. Cells were pelleted, washed with 300 μL of 0.1 M LiAc twice, then pelleted and resuspended in 30 μL of 0.1 M LiAc. Then, with the cells on ice, the following were then added to the cells in order: 100 μL 50% w/v PEG 3350, 15 μL 1.0 M LiAc, 6 μL 7 mg/mL ss carrier DNA, 18 μL DMSO, 15 μL PCR product, and 14 μL sterile water. This was mixed and incubated in a GeneAMP™ PCR System 9700 (Thermofisher Scientific) for 30 minutes at 30˚C, followed by 15 minutes at 42˚C. Cells were pelleted, the transformation buffer was then removed by aspiration, and cells were resuspended in 100 μL water pre-warmed to 30˚C. All 100 μL of cells were then spread on an agar plate containing the appropriate selection. For histidine selection, plates were made from synthetic complete media lacking histidine. For G418 selection, cells were first plated on a YPD plate and then replica plated to a YPD plate including 600 μg/mL G418 the next morning. Plates were incubated at 30˚C for 2-3 days. A full list of strains used in this study are listed in Supplementary Table 2. Insertion of the desired construct in the genome was checked via colony PCR.

**Preparing cells for microscopy**

To prepare cells for microscopy, a single colony was picked from an agar plate into 500 μL synthetic complete media. 100 μL of this cell suspension was then pipetted into 400 uL synthetic complete media, to obtain a 1:5 dilution as well. The cells were grown for 16-20 hrs in a shaking incubator at 30 ˚C. Whichever culture had an approximate $OD_{600}$ of 0.1-0.8 was then used for imaging.

To prepare slides for imaging, 22x40 mm glass coverslips with thickness no. 1 (VWR) were cleaned using a 40-minute exposure to argon plasma in a 2.6 L Zepto plasma laboratory unit (Diener Electronic). Frame-Seal slide chambers (9 × 9 mm2, Biorad, Hercules, CA) were attached to the plasma-cleaned coverslips and the surface treated with 100 µL concanavalin A (Sigma-Aldrich) (2 mg/mL in PBS). The concanavalin A was removed from the surface after 30 s by aspirating with a pipette. 100 µL of prepared yeast culture was then pipetted onto the slide and left for approximately 5 minutes to allow cells to attach. The cells were then aspirated from the slide and the surface washed using 100 µL fresh PBS and gently pipetting up and down. 100 µL fresh PBS was then added to the slide before imaging.

**TIRF microscopy**

Microscopy data was primarily collected using a commercial TIRF microscope (Oxford Nanoimager). Images were acquired with an exposure of 50 ms for between 2000 and 4000 frames using the NimOS software. mNeonGreen was activated using the 488 nm laser at 15% power and the angle of illumination (TIR angle) was set at 48.5° for all acquisitions. Pixel length was 117 nm. All images were collected at room temperature.

Two color, Cdc12 and Pil1 microscopy data was collected using on a custom built TIRF microscope. mNG was excited using a 488 nm laser (Cobolt MLD 488-200 Diode Laser System, Cobalt, Sweden) ~40W/cm$^2$ and mCherry was excited using a 561 nm laser (Cobolt DPL Series 561-100 DPSS Laser System, Cobolt, Sweden) ~25W/cm$^2$. Both lasers were aligned and directed parallel to the optical axis at the edge of a 1.49 NA TIRF objective (CFI Apochromat TIRF 60XC Oil, Nikon, Japan), mounted on an inverted Nikon TI2 microscope (Nikon, Japan). To prevent z-stage drift during imaging a perfect focus system was utilized. Fluorescence collected by the same objective was separated from the returning TIR beam by a dichroic mirror (Di01-R405/488/561/635 (Semrock, Rochester, NY, USA)), and was passed through appropriate filters (488 nm: BLP01-488R, FF01-520/44 (Semrock, NY, USA), 561 nm: LP02-568- RS, FF01-587/35 (Semrock, NY, USA)). Fluorescence was then passed through a 2.5× beam expander and recorded on an EMCCD camera (Delta Evolve 512, Photometrics) operating in frame transfer mode (EMGain = 11.5 e- /ADU and 250 ADU/photon). Each pixel was 103 nm in length. The microscope was automated by using the open-source microscopy platform Micromanager. For two color samples, images were recorded by collecting 200 frames of 50 ms using the 561 nm laser followed by 200 frames of 50 ms using the 488 nm laser. This was repeated 10 times to result in a total of 2,000 frames for both mCherry and mNeonGreen. All Cdc12 and Pil1 images, were acquired using the 488 nm laser using an exposure of 50 ms for between 2000 and 4000 frames. All images were collected at room temperature.

**Confocal microscopy**

For localization of Pma1 throughout entire cells, a Zeiss LSM880 confocal microscope (with alpha Plan-Apochromat 100x/1.46 Oil DIC M27 Elyra objective) and Airyscan was used to capture mNG fluorescence data (ex/em 488/522) with z-stack increments of 0.170 micron. Airyscan image processing for generation of super-resolution 3D data was carried out in Zen Blue (Zeiss).

**Super-resolution analysis**

Super-resolution images were analyzed using Fiji (Java 8 2017 release). Single localizations were processed using the Peak Fit function of the Fiji GDSC SMLM plugin, using a signal strength threshold of 30, a minimum photon threshold of 100, and a precision threshold of 15-30 nm. All localization files are available at: https://doi.org/10.5281/zenodo.8101098. The overall resolution of the images was calculated using Fourier Ring Correlation analysis using the Python code available at: https://doi.org/10.5281/zenodo.7275952.

**Quantification of membrane specific localizations**

First, cell segmentation on the maximum intensity projections was performed using an adaptive Otsu algorithm on CellProfiler.[5] Segmented cell objects were shrunk down by 4 pixels and subtracted from the cell mask to generate membrane masks. The code available at https://doi.org/10.5281/zenodo.7817411 was then used to tag localizations contained within localization files generated through super-resolution analysis described previously as cell, membrane or external based on overlap with the respective masks. Cells with fewer than 50 localizations were excluded from further analysis.

**Cluster analysis**

DBSCAN[6] complied in Python 3.8 using epsilon = 0.8 pixels and a minimum points threshold of 3 was used to identify clusters of localizations in the localization files generated through super-resolution analysis described previously. For cells with more than 10 clusters, several parameters were then calculated and recorded: the number of localizations within each cluster, the ID of the nearest cluster, and the distance to the nearest cluster (determined using unsupervised nearest neighbors learning with the Ball Tree algorithm). This information was collated for all images of cells with the same protein target and peptide pair used for labelling. The Python code is available at: https://doi.org/10.5281/zenodo.8060555. For two color images, cluster analysis was carried out on localization data for each color as described previously. The distance between each mCherry (Pil1) cluster and the nearest mNeonGreen (Arc35) cluster in the same cell was determined by calculating the Euclidian distance to all the mNG clusters in the cell and recording the shortest distance. This information was collated for all two-color images of Arc35 and Pil1. The Python code is available at: https://doi.org/10.5281/zenodo.8060654.

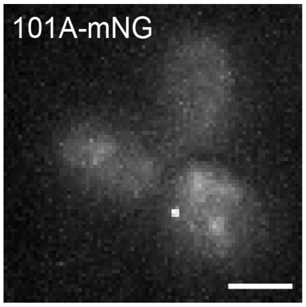


**Figure S1**. 101A-mNeonGreen alone does not localize to the plasma membrane. Diffraction-limited TIRF image of yeast expressing 101A-mNG but no protein tagged with 101B. No appreciable membrane localization is visible. Scale bar is 2 μm.

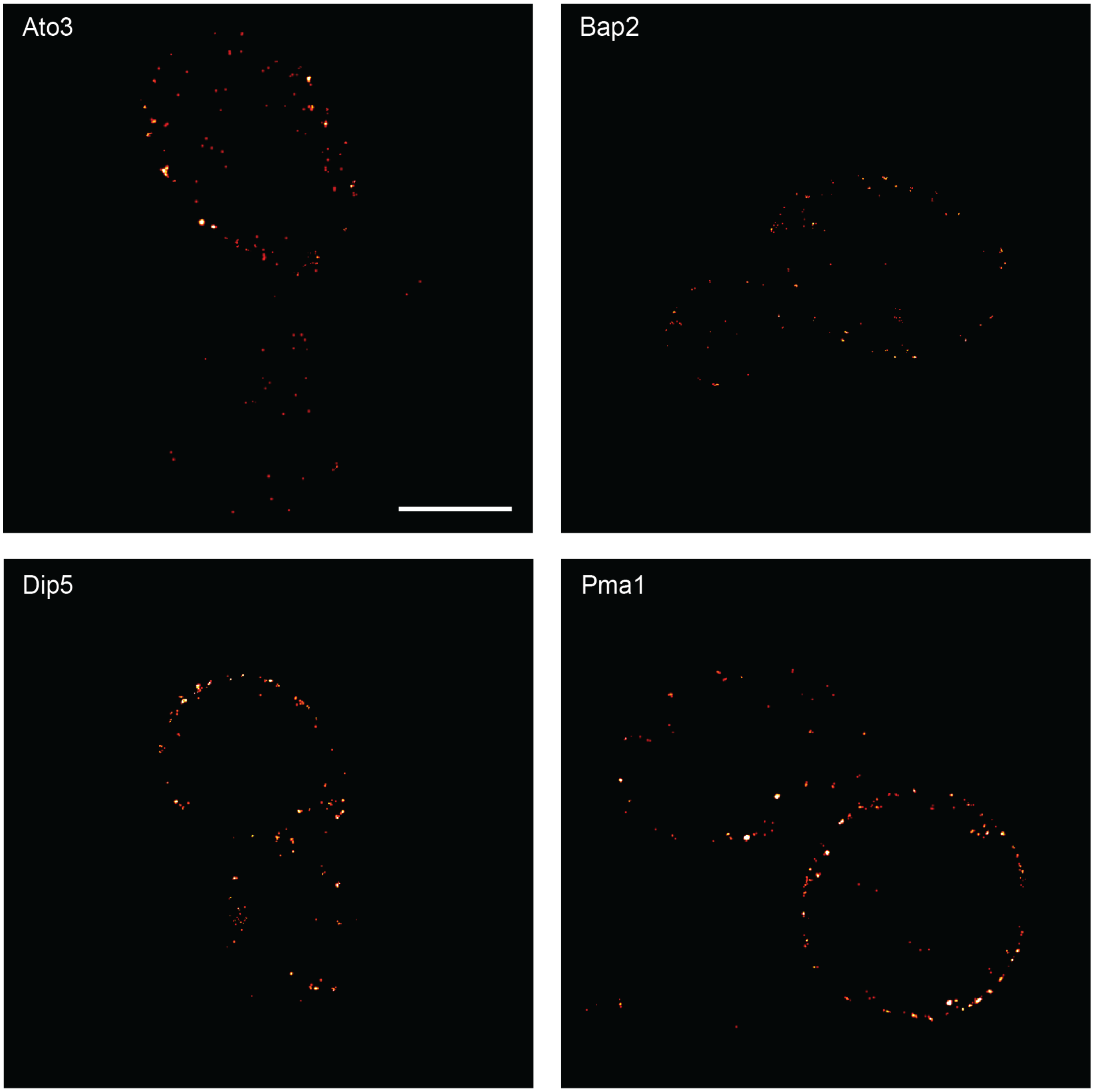


**Figure S2**. Full-cell super-resolution images of membrane proteins labelled with the 101A/B peptide pair shown in Figure 3. Scale bar is 2 μm and all images have the same dimensions.

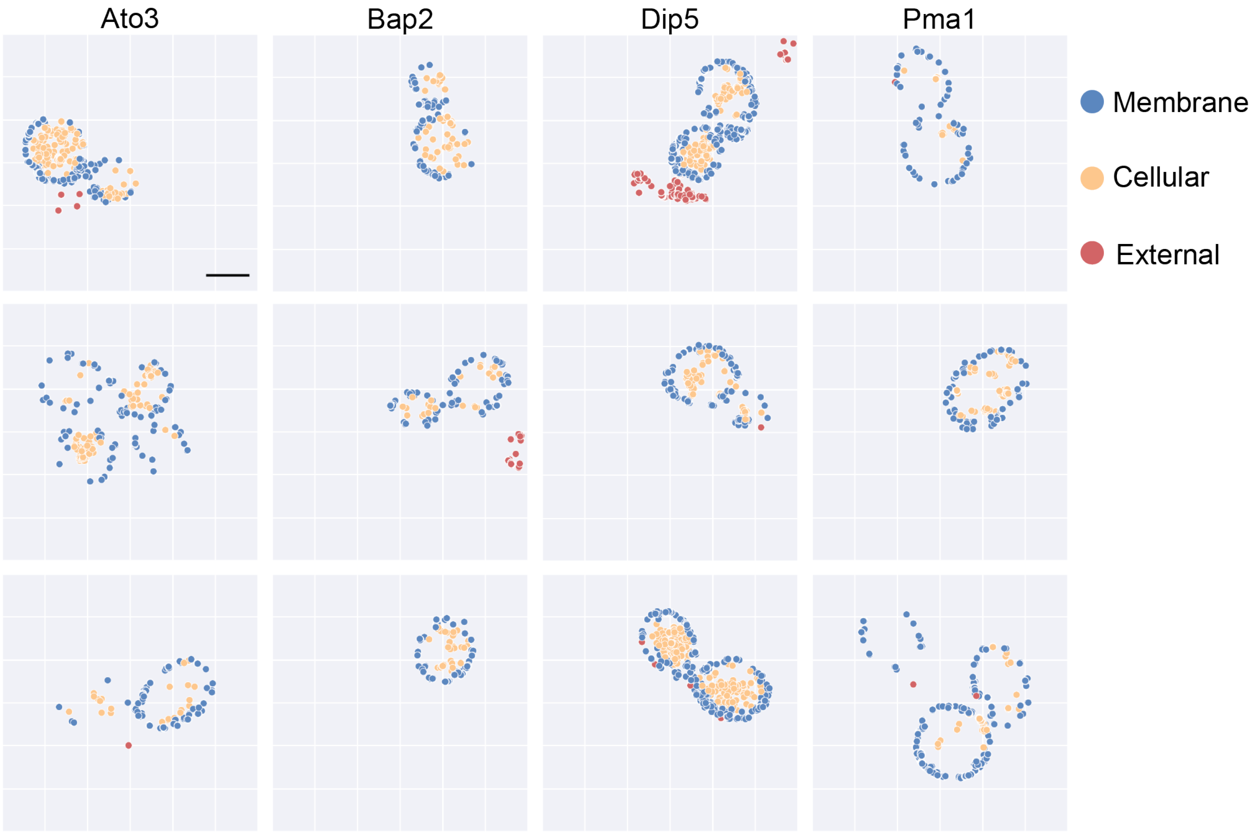


**Figure S3.** Representative plots showing localizations classified as membrane, cellular or external for membrane proteins labelled with the 101A/B peptide pair. Localizations classed as membrane are plotted in blue, cellular in yellow and external in red. Scale bar is 2 µm and all images have the same dimensions.

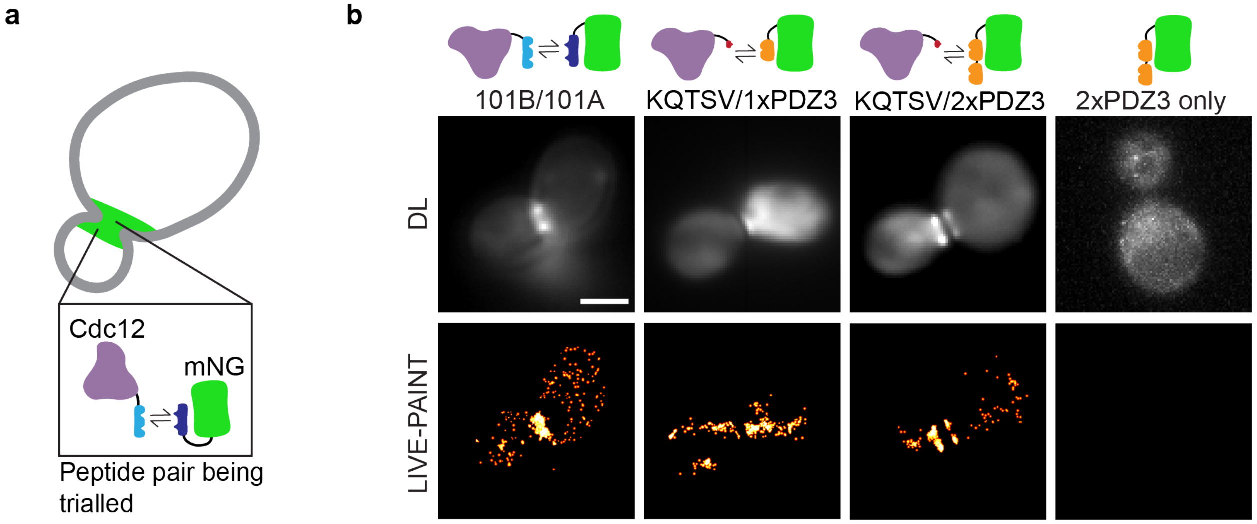


**Figure S4.** KQTSV can be used as a peptide label for LIVE-PAINT with two tandem repeats of PDZ3 fused to mNeonGreen. (a) Schematic representation of the localization of the septum protein Cdc12 in a dividing *S. Cerevisiae* cell and the LIVE-PAINT labelling strategy used to image this protein. (b) Diffraction-limited (DL) (top) and LIVE-PAINT (bottom) images of Cdc12 imaged by C-terminal labelling with 101B and co-expressing 101A-mNG or by C-terminal labelling with KQTSV and either co-expressing 1xPDZ3-mNG or 2xPDZ3-mNG and control images of 2xPDZ3-mNG only. Scale bar is 2 µm and all images have the same dimensions.

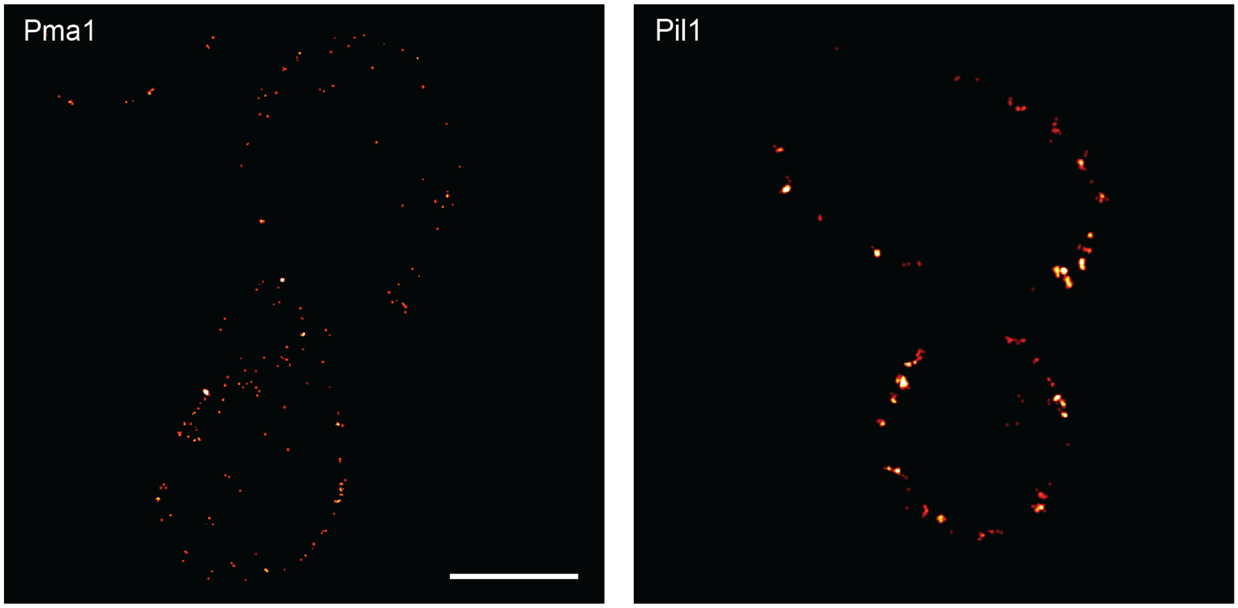

**Figure S5**. Full-cell super-resolution images of membrane proteins labelled with the KQTSV/2xPDZ3 peptide-protein pair shown in Figure 4. Scale bar is 2 μm and all images have the same dimensions.

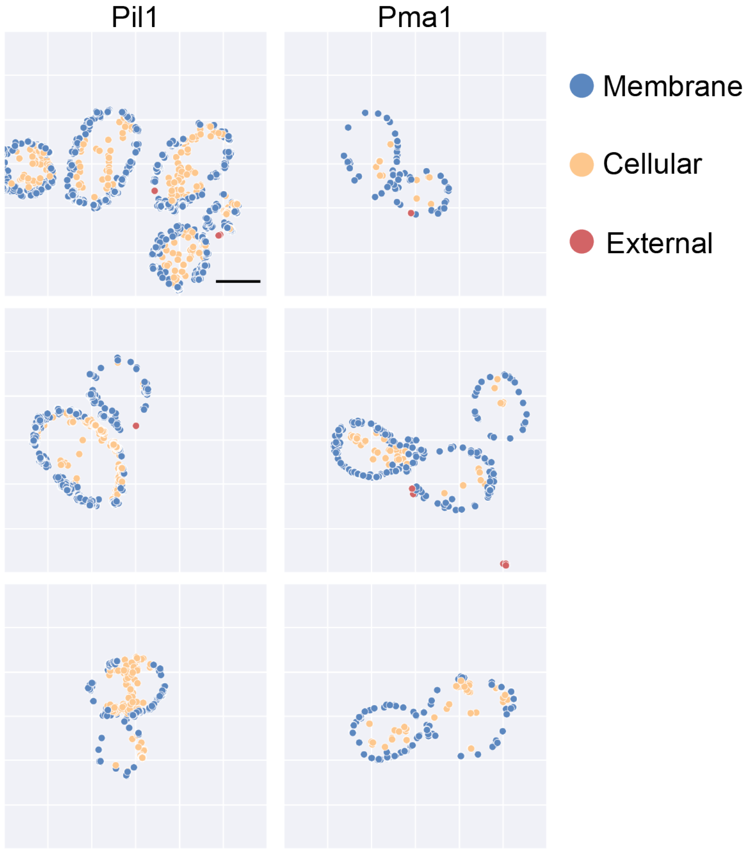


**Figure S6.** Representative plots showing localizations classified as membrane, cellular or external for membrane proteins labelled with the KQTSV/2xPDZ3 peptide-protein pair. Localizations classed as membrane are plotted in blue, cellular in yellow and external in red. Scale bar is 2 μm and all images have the same dimensions.

**Table S1.** Membrane transporter proteins tagged and imaged in this study.

| **Protein** | **Number of proteins/cell**[7] | **Protein function**[7] |
|---|---|---|
| Ady2 | 1,298 | Transporter protein required for ammonia export and acetate uptake and resistance. |
| Agp1 | 4,012 | Broad substrate range permease that transports asparagine and glutamine with intermediate specificity. |
| Ato3 | 2,211 | Transporter protein required for ammonia export. |
| Bap2 | 6,997 | Permease for leucine, valine and isoleucine. Also transports cysteine, methionine, phenylalanine, tyrosine and tryptophan. |
| Can1 | 3,618 | High-affinity permease for arginine. |
| Dip5 | 5,961 | Can transport glutamate, aspartate, glutamine, asparagine, serine, alanine and glycine. |
| Fcy2 | 5,492 | This permease has a broad specificity towards purines, and also transport cytosine and 5-methylcytosine but neither uracil nor thymine. |
| Gap1 | 3,634 | General amino-acid permease involved in the uptake of all the naturally occurring L-amino-acids, related compounds such as ornithine and citrulline, some D-amino acids, toxic amino acid analogs such as azetidine-2-carboxylate, and the polyamines putrescine and spermidine. |
| Mup1 | 4,102 | High affinity permease for methionine. |
| Pma1 | 40,732 | The plasma membrane ATPase of plants and fungi is a hydrogen ion pump. |
| Ptr2 | 2,551 | Uptake of small peptides. |
| Sit1 | 2,822 | Involved in the transport of siderophore ferrioxamine B and so has a role in iron homeostasis. |

**Table S2.** Summary statistics of super-resolution images of membrane associated proteins imaged using LIVE-PAINT in Figure 3.

| Protein | Number of Cells | Average Resolution (nm) | Average Precision (nm) | Average Number of Localisations |
|---|---|---|---|---|
| Pma1 | 15 | 67.28 +/- 13.40 | 10.73 +/- 0.40 | 367 +/- 315 |
| Bap2 | 15 | 61.40 +/- 15.03 | 10.56 +/- 1.00 | 207 +/- 95 |
| Dip5 | 6 | 84.44 +/- 20.53 | 10.32 +/- 0.44 | 474 +/- 294 |
| Ato3 | 6 | 73.46 +/- 24.55 | 10.69 +/- 1.18 | 492 +/- 565 |

Resolution was calculated using FRC and all values are shown as mean +/- SD.

**Table S3.** Summary statistics of super-resolution images of membrane associated proteins imaged using LIVE-PAINT and the peptide-protein pair KQTSV/2xPDZ3 in Figure 4.

| Protein | Number of Cells | Average Resolution (nm) | Average Precision (nm) | Average Number of Localisations |
|---|---|---|---|---|
| Pma1 | 5 | 113.61 +/- 52.70 | 10.93 +/- 0.21 | 204 +/- 104 |
| Pil1 | 5 | 101.19 +/- 17.12 | 20.5 +/- 5.88 | 1157 +/- 718 |

Resolution was calculated using FRC and all values are shown as mean +/- SD.

**Table S4.** Summary statistics of super-resolution images of Arc35 and Pil1 imaged using two color LIVE-PAINT and shown in Figure 5.

| Protein | Fluorescent protein | Average Resolution (nm) | Average Precision (nm) | Average Number of Localisations |
|---|---|---|---|---|
| Arc35 | mNG | 83.03 +/- 21.51 | 15.59 +/- 3.61 | 657 +/- 475 |
| Pil1 | mCherry | 72.45 +/- 27.94 | 19.73 +/- 2.87 | 620 +/- 380 |

Resolution was calculated using FRC and all values are shown as mean +/- SD. Number of cells = 6.

**Table S5.** Primers for tagging membrane proteins at their genomic loci.

| Name | Sequence |
|---|---|
| p6h_int_F | CTAATCCAAGGAGGTTTACGGACCAGGGGAACTTTCCAGATTCAG AAGCTTCGTACGCTGCA |
| p6h_int_R | CATGAAAAATTAAGAGAGATGATGGAGCGTCTCACTTCAAACGCA GGCGTTAGTATCGAATCG |
| ADY2_F | TCGTCCATTCCCATTACCATCTACTGAAAGGGTAATCTTTGGTGGA TCAGGCTCTGG |
| ADY2_R | TTTTTATTTCAATAGTTCTCGTTATTAGTAGGTCGTGCTCATTAGAA AAACTCATCGAGCATC |
| AGP1_F | GAACGGACCTTATTGGAAAAGGGTCGTTGCCTTCTGGTGTGGTGGA TCAGGCTCTGG |
| AGP1_R | CAAAAATGAATAAATATAAAAGAAGTAAATGCTTTTTTTATTAGA AAAACTCATCGAGCATC |
| ATO3_F | TTATTTAGCCTTCAGGGCGCACACAATGCCAAATGCTCCTGGTGGA TCAGGCTCTGG |
| ATO3_R | TTTTAATGTTTTATAAGTTTTGTTTTTCATTTCATACCCTATTAGAAA AACTCATCGAGCATC |
| BAP2_F | GAATATGTCTTTGATGAGAAAAGCTTATCATTTCTGGTGTGGTGGA TCAGGCTCTGG |
| BAP2_R | TCTAATGGGAAGTGTCCAGACCTGAGTGGTGTAGTTAAGTATTAGA AAAACTCATCGAGCATC |
| CAN1_F | ACCAAAGACTTTTTGGGACAAATTTTGGAATGTTGTAGCAGGTGGA TCAGGCTCTG |
| CAN1_R | ATGGCGTGGAAATGTGATCAAAGGTAATAAAACGTCATATATGGC GGCGTTAGTATC |
| DIP5_F | TATGGAGTGGTTCTATGAAAAATTTTTGGGTAATATCTTCGGTGGA TCAGGCTCTG |
| DIP5_R | ATAGTTTCATGTTGCCCTAAGGCTTTGATTAAAAAGGCATATGGCG GCGTTAGTATC |

| | |
|---|---|
| FCY2_F | CAACATCTTAAGACCTTTAGAATTAAAATACTTCGGTCGTGGTGGA TCAGGCTCTG |
| FCY2_R | GAAATGTGCACGGGGAAATGATCGCCCTAATCATTACTTCATGGCG GCGTTAGTATC |
| GAP1_F | CACAAAGCCAAGATGGTATAGAATCTGGAATTTCTGGTGTGGTGG ATCAGGCTCTG |
| GAP1_R | ATCTAAAAAATAAAGTCTTTTTTTGTCGTTGTTCGATTCAATGGCGG CGTTAGTATC |
| MUP1_F | TATAATCGAACATTACAAAAGTGAACAAGAAAAATCGCTGGGTGG ATCAGGCTCTG |
| MUP1_R | GATTATAAGAATCGAGATGAGATGGTAAGTACCTTTTTGGATGGCG GCGTTAGTATC |
| PMA1_F | TGCTATGCAAAGAGTCTCTACTCAACACGAAAAGGAAACCGGTGG ATCAGGCTCTG |
| PMA1_R | AATGTGACAAAAATTATGATTAAATGCTACTTCAACAGGAATGGC GGCGTTAGTATC |
| PMA1_URA3_R | AATGTGACAAAAATTATGATTAAATGCTACTTCAACAGGATCAGTT TTGCTGGCCGC |
| PTR2_F | ATTAGAACCAATGGAAAGTCTAAGATCCACCACCAAATATGGTGG ATCAGGCTCTG |
| PTR2_R | AAAAAAAAAAAAAAAAAAAGACAGTAAGTTAATTAAACGCAATGG CGGCGTTAGTATC |
| SIT1_F | ATTCTTTACGCACTTTACAAGCAGTAAAGATAGGAAAGATGGTGG ATCAGGCTCTG |
| SIT1_R | GCTATATGTGCATGTATGAAATTATTTGGGTGAGATAATAATGGCG GCGTTAGTATC |

p6h_int_F and p6h_int_R were used to amplify the plasmid containing pGAL1 101A-mNG, pGAL1 PDZ3-mNG, or pGAL1 2xPDZ3-mNG and HIS3 selection marker and to integrate it into the genome replacing GAL2. PMA1_URA3_R was used with PMA1_F to amplify the plasmid containing the KQTSV peptide and the URA3 selection marker. The ADY2, AGP1, ATO3, and BAP3 primers were used to amplify the plasmid containing the 101B peptide and the G418 selection marker. All other primers were used to amplify the plasmid containing the 101B peptide and the URA3 selection marker. The primer name contains the name of the gene being tagged and the primers come in pairs, with the "_F" and "_R" primers being used as forward and reverse primers.

**Table S6**. Primers for generating a yeast strain with two membrane-associated proteins tagged at their genomic loci.

| Name | Sequence |
|---|---|
| ARC35_F | ACAGGCAAGAAGAACCTTCACCGGTAGAAAGATTGTCTACGGTGGATCAGGCTCTGG |
| ARC35_R | TAACCCTTTTTACGGATTCTTACGTACTTATTTAATCTTTATTAGAAAAACTCATCGAGCATC |
| PIL1_MULTI_F | ACACCAGCAAAGTGAGTCTCTTCCCCAACAAACAACAGCTGGTGGATCAGGCTCTGG |
| PIL1_MULTI_R | TTTTTTTTTTTGTTTCTAATAGATTGTTGATTTATTTTGAGGCGTTAGTATCGAATCGAC |

The primers come in pairs, with the "_F" and "_R" primers being used as forward and reverse primers. Primers ARC35_F/R were used to fuse 101B to Arc35 by amplifying the plasmid containing 101B and the G418 selection marker. PIL1_MULTI_F/R were used to fuse 108B to PIL1 and insert pGAL1 101A-mCherry at the same gentic loci by amplifying the plasmid containing 108B, pGAL1-108A-mCherry and the selection marker LEU2.

**Table S7.** List of yeast strains used in this study.

| # | Gene | Coiled-coil pair | Genotype |
|---|---|---|---|
| 1 | ADY2 | 101A/B | MATa his3Δ1 leu2Δ0 met15Δ0 ura3Δ0 gal2Δ::HIS3MX6 pGAL1 101A-mNeonGreen ADY2-101B::KANMX6 |
| 2 | AGP1 | 101A/B | MATa his3Δ1 leu2Δ0 met15Δ0 ura3Δ0 gal2Δ::HIS3MX6 pGAL1 101A-mNeonGreen AGP1-101B::KANMX6 |
| 3 | ATO3 | 101A/B | MATa his3Δ1 leu2Δ0 met15Δ0 ura3Δ0 gal2Δ::HIS3MX6 pGAL1 101A-mNeonGreen ATO3-101B::KANMX6 |
| 4 | BAP2 | 101A/B | MATa his3Δ1 leu2Δ0 met15Δ0 ura3Δ0 gal2Δ::HIS3MX6 pGAL1 101A-mNeonGreen BAP2-101B::KANMX6 |

| | | | |
|---|---|---|---|
| 5 | CAN1 | 101A/B | MATa his3Δ1 leu2Δ0 met15Δ0 ura3Δ0 gal2Δ::HIS3MX6 pGAL1 101A-mNeonGreen CAN1-101B::URA3 |
| 6 | DIP5 | 101A/B | MATa his3Δ1 leu2Δ0 met15Δ0 ura3Δ0 gal2Δ::HIS3MX6 pGAL1 101A-mNeonGreen DIP5-101B::URA3 |
| 7 | FCY2 | 101A/B | MATa his3Δ1 leu2Δ0 met15Δ0 ura3Δ0 gal2Δ::HIS3MX6 pGAL1 101A-mNeonGreen FCY2-101B::URA3 |
| 8 | GAP1 | 101A/B | MATa his3Δ1 leu2Δ0 met15Δ0 ura3Δ0 gal2Δ::HIS3MX6 pGAL1 101A-mNeonGreen GAP1-101B::URA3 |
| 9 | MUP1 | 101A/B | MATa his3Δ1 leu2Δ0 met15Δ0 ura3Δ0 gal2Δ::HIS3MX6 pGAL1 101A-mNeonGreen MUP1-101B::URA3 |
| 10 | PMA1 | 101A/B | MATa his3Δ1 leu2Δ0 met15Δ0 ura3Δ0 gal2Δ::HIS3MX6 pGAL1 101A-mNeonGreen PMA1-101B::URA3 |
| 11 | PTR2 | 101A/B | MATa his3Δ1 leu2Δ0 met15Δ0 ura3Δ0 gal2Δ::HIS3MX6 pGAL1 101A-mNeonGreen PTR2-101B::URA3 |
| 12 | SIT1 | 101A/B | MATa his3Δ1 leu2Δ0 met15Δ0 ura3Δ0 gal2Δ::HIS3MX6 pGAL1 101A-mNeonGreen SIT1-101B::URA3 |
| 13 | PMA1 | 2xPDZ3/ KQTSV | MATa his3Δ1 leu2Δ0 met15Δ0 ura3Δ0 gal2Δ::HIS3MX6 pGAL1 PDZ3-mNeonGreen PMA1-KQTSV::URA3 |
| 14 | PIL1 | 2xPDZ3/ KQTSV | MATa his3Δ1 leu2Δ0 met15Δ0 ura3Δ0 gal2Δ::HIS3MX6 pGAL1 PDZ3-mNeonGreen PIL1-KQTSV::URA3 |
| 15 | ARC35 & PIL1 | 101A/B and 108A/B | MATa his3Δ1 leu2Δ0 met15Δ0 ura3Δ0 gal2Δ::HIS3MX6 pGAL1 101A-mNeonGreen ARC35-101B::KANMX6 Pil1-108B pGAL1 108A-mCherry::LEU2 |

The gene being tagged is listed in the "Gene" column and the full genotype of the strain used for imaging is given in the "Genotype" column.

**References**


(1) Guthrie, C.; Fink, G. R. *Guide to Yeast Genetics and Molecular and Cell Biology, Part C*; Gulf Professional Publishing, 2002.

(2) Thompson, K. E.; Bashor, C. J.; Lim, W. A.; Keating, A. E. SYNZIP Protein Interaction Toolbox: In Vitro and in Vivo Specifications of Heterospecific Coiled-Coil Interaction Domains. *ACS Synth. Biol.* **2012**, *1* (4), 118–129. https://doi.org/10.1021/sb200015u.

(3) Oi, C.; Gidden, Z.; Holyoake, L.; Kantelberg, O.; Mochrie, S.; Horrocks, M. H.; Regan, L. LIVE-PAINT Allows Super-Resolution Microscopy inside Living Cells Using Reversible Peptide-Protein Interactions. *Commun. Biol.* **2020**, *3* (1), 1–10. https://doi.org/10.1038/s42003-020-01188-6.

(4) Hawkins, K. M.; Smolke, C. D. The Regulatory Roles of the Galactose Permease and Kinase in the Induction Response of the GAL Network in Saccharomyces Cerevisiae. *J. Biol. Chem.* **2006**, *281* (19), 13485–13492. https://doi.org/10.1074/jbc.M512317200.

(5) Stirling, D. R.; Swain-Bowden, M. J.; Lucas, A. M.; Carpenter, A. E.; Cimini, B. A.; Goodman, A. CellProfiler 4: Improvements in Speed, Utility and Usability. *BMC Bioinformatics* **2021**, *22* (1), 433. https://doi.org/10.1186/s12859-021-04344-9.

(6) Sander, J.; Ester, M.; Kriegel, H.-P.; Xu, X. Density-Based Clustering in Spatial Databases: The Algorithm GDBSCAN and Its Applications. *Data Min. Knowl. Discov.* **1998**, *2* (2), 169–194. https://doi.org/10.1023/A:1009745219419.

(7) Cherry, J. M.; Hong, E. L.; Amundsen, C.; Balakrishnan, R.; Binkley, G.; Chan, E. T.; Christie, K. R.; Costanzo, M. C.; Dwight, S. S.; Engel, S. R.; Fisk, D. G.; Hirschman, J. E.; Hitz,

B. C.; Karra, K.; Krieger, C. J.; Miyasato, S. R.; Nash, R. S.; Park, J.; Skrzypek, M. S.; Simison, M.; Weng, S.; Wong, E. D. Saccharomyces Genome Database: The Genomics Resource of Budding Yeast. *Nucleic Acids Res.* **2012**, *40* (Database issue), D700. https://doi.org/10.1093/nar/gkr1029.
